# Enterovirus A119 in a child with Acute Flaccid Paralysis, Nigeria

**DOI:** 10.1101/084566

**Authors:** JA Adeniji, AO Oragwa, UE George, UI Ibok, TOC Faleye, MO Adewumi

## Abstract

The oldest EV-A119 record was in 2008 in a chimpanzee in Cameroon and subsequently in more non-human primates and healthy children. Here we report for the first time the detection of EV-A119 in a child with Acute Flaccid Paralysis, thus suggesting possible association with a clinical condition in humans.

## Short Communication

Enteroviruses are members of the genus *Enterovirus* in the family *Picornaviridae*, order *Picornavirales*. The type member of the genus is poliovirus, the etiologic agent of poliomyelitis. In 1988, the WHA resolved to eradicate poliovirus, and as part of its strategy, intensive surveillance is carried out among children <15 years old with Acute flaccid paralysis (AFP). In Nigeria, when surveillance and laboratory detection as recommended by the WHO cell culture based algorithm (WHO, 2004) are optimal, a minimum of 10% of stool from all AFP cases contain enteroviruses. However, we (Adeniji and Faleye, 2015) have previously shown that the cell culture based algorithm (WHO, 2004) biases our perception of the enterovirus diversity of a sample by not fully detecting all the types present therein. This therefore supposes that several samples declared negative might contain enterovirus types for which the cell lines in the algorithm might not be susceptible and permissive.

In this light, RNA was extracted from 30 archived stool suspensions collected in 2015 from children <15 years old diagnosed with AFP. The samples were collected in accordance with national ethical guidelines as part of the national poliovirus surveillance programme and sent to the WHO National Polio Laboratory in Ibadan, Nigeria to screen for the likely presence of poliovirus. The samples were consequently considered negative because they showed no cytopathology in both RD and L20B cell lines (WHO, 2004). These samples were then anonymized and 30 of such randomly selected for this study. Using the RNA extract, cDNA synthesis and RT-semi-nested PCR (RT-snPCR) amplifying an ~350bp stretch in 5’ end of the VP1 gene was done as recently recommended by the WHO for enterovirus surveillance (WHO, 2015), and previously described (Faleye et al., 2016). One of the amplicons sequenced was identified by the enterovirus genotyping tool (Kroneman et al., 2011) as EV-A119 (GenBank accession number: KX765171). To confirm this observation, the assay was repeated but instead of using primer AN89 as the forward primer for the second round PCR assay, primer 189 which has a predilection for enterovirus species A and C was used (WHO, 2015). The sequenced amplicon again confirmed the enterovirus present as EV-A119.

The earliest record of EV-A119 in GenBank was that detected in a Chimpanzee in Cameroon in 2008 (Sadeuh-Mba et al., 2014). In 2009, it was detected in a healthy child in the same country (Ayukekbong et al., 2013). By 2010, it was again detected in both a Chimp and a Gorilla in Cameroon (Harvala et al., 2014). In 2013, it was detected in a healthy child in Cote d’Ivoire (Di Cristianzano, et al., 2015) (Figure 1). Against this backdrop, finding for the first time to the best of our knowledge, EV-A119 in a child diagnosed with AFP in Kaduna, Nigeria is worthy of report. Though it might not be definitive evidence of EV-A119s role in the AFP, it is course for concern because it raises the possibility of such. Also, it might be suggestive of the likelihood that consequent to the previous “uneventful” spill-over events (Ayukekbong et al., 2013; Di Cristianzano, et al., 2015), the virus might have now become well adapted to humans and pathogenic strains evolved. Furthermore, the phylogenetic evidence might be suggestive that several strains of EV-A119 are circulating undetected (Figure 1).

**Figure 1:**
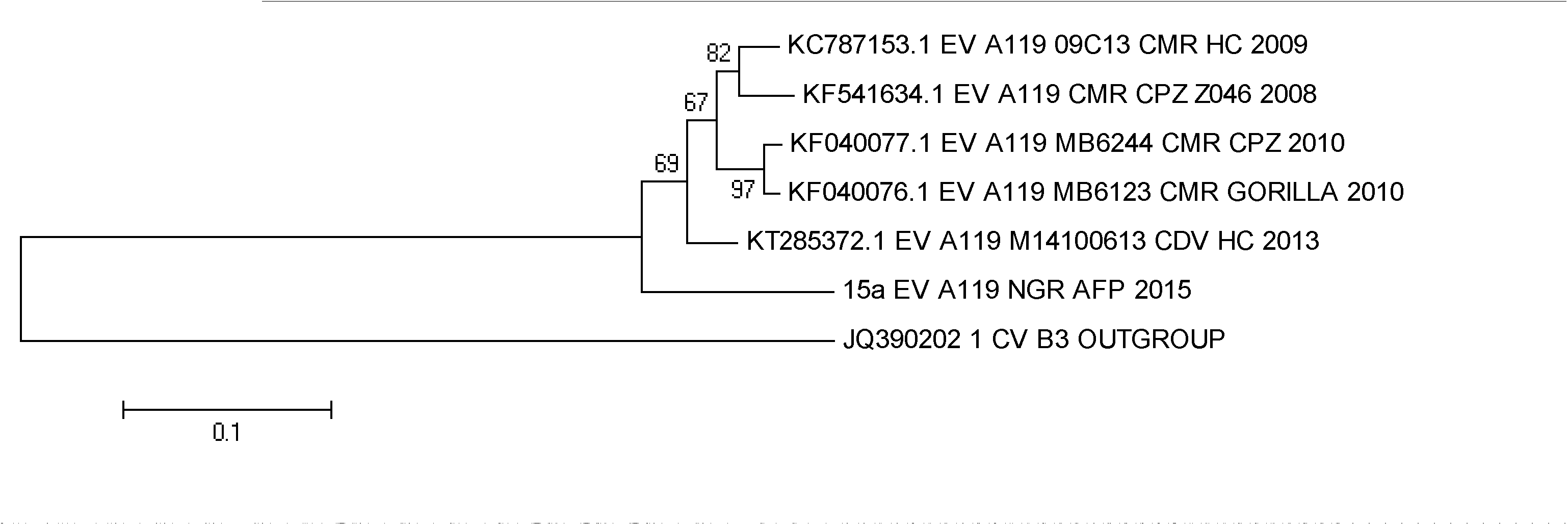
Phylogeny of all EV-A119 strains publicly available in GenBank as at the 23^rd^ of August 2016 and the isolate (15a) described in this study.

To the best of our knowledge, no group has reported EV-A119 growing in any cell line and in this study, it was also detected in a stool suspension that was negative for enteroviruses by cell culture. This implies that the WHO cell culture based algorithm (WHO, 2004) failed to detect it. It is noteworthy that a good number of enteroviruses recently detected and first described in non-human primates (NHPs) could not be detected using cell culture based algorithms and were therefore detected using the RT-snPCR algorithm used in this study (Harvala et al., 2014; Oberste et al., 2013; Sadeuh-Mba et al., 2014). The question then is how many more enterovirus types are out there already adapting or adapted to humans but yet to be described because of the limit and biases of the cell culture based algorithm. Therefore, the need to effectively and efficiently search for all enteroviruses previously documented in NHPs (Harvala et al., 2014; Oberste et al., 2013; Sadeuh-Mba et al., 2014) cannot be overemphasized. How long EV-A119 has been spilling into human populations is currently not clear but its zoonotic potential is unmistakeable. Considering the epidemiology of poliovirus and other enteroviruses, the scope and geographical spread of this enterovirus type, that has till date only been detected in sub-Saharan Africa, and might be associated with AFP, deserves to be defined.

## ACKNOWLEDGEMENTS

We thank the WHO National Polio Laboratory in Ibadan, Nigeria for providing the anonymous samples analyzed in this study. This study was funded by contributions from the authors.

